# A multiple comorbidities mouse model to assess atherosclerosis progression following lung infection in *ApoE* deficient mice

**DOI:** 10.1101/2022.03.23.485412

**Authors:** Benjamin Bartlett, Silvia Lee, Herbert P Ludewick, Teck Siew, Shipra Verma, Grant Waterer, Vicente F. Corrales-Medina, Girish Dwivedi

**Affiliations:** Department of Advanced Clinical and Translational Cardiovascular Imaging, Harry Perkins Institute of Medical Research, Murdoch, Western Australia; School of Medicine, University of Western Australia, Australia; Department of Cardiology, Fiona Stanley Hospital, Murdoch, Western Australia, Australia; Department of Microbiology, Pathwest Laboratory Medicine, Perth, Australia; Department of Nuclear Medicine, Fiona Stanley Hospital, Murdoch, Western Australia; Department of Geriatric Medicine, Fiona Stanley Hospital, Murdoch, Western Australia; Heart and Lung Research Institute, Harry Perkins Institute of Medical Research, Murdoch, Western Australia, Australia; Department of Nuclear Medicine, Sir Charles Gairdner Hospital, Western Australia, Australia; Royal Perth Hospital, Victoria Square, Perth, Western Australia, Australia; Department of Medicine, University of Ottawa, Canada; Clinical Epidemiology Program, The Ottawa Hospital Research Institute, Ottawa, Canada

**Author notes:** Corresponding author: Professor Girish Dwivedi, MD, PhD, MRCP, FASE, FESC, FRACP, FSCCT, FCSANZ Wesfarmers Chair in Cardiology and Consultant Cardiologist, Harry Perkins Institute of Medical Research and Fiona Stanley Hospital, The University of Western Australia, Perth, Australia.

**Keywords:** Atherosclerosis, ApoE deficient mice, Pneumonia

## Abstract

**Background:** Inflammation is a risk factor for atherosclerosis progression. Hospitalisation for pneumonia is associated with increased risk of cardiovascular disease. Herein, we describe a multiple comorbidities murine model to study the impact of bacterial pneumonia on atherosclerosis.

**Methods:** Firstly, a minimal infectious dose of *Streptococcus pneumoniae* (TIGR4 strain) to produce clinical pneumonia with a low mortality rate (20%) was established. C57Bl/6 *ApoE-/-* mice were fed a high-fat diet prior to administering intranasally 10^5^ colony forming units of TIGR4 or phosphate buffered saline (PBS). At days 2, 7 and 28 post inoculation (PI), the lungs of mice were imaged by MRI and PET. Mice were euthanised and investigated for changes in systemic inflammation and changes in lung morphology using ELISA, Luminex assay and real-time PCR.

**Results:** TIGR4 inoculated mice presented with varying degreess of lung infiltrate, pleural effusion and consolidation on MRI at all timepoints up to 28 days PI. Moreover, PET scans identified significantly higher FDG uptake in the lungs of TIGR4 inoculated mice up to 28 days PI. Majority (90%) TIGR4-inoculated mice developed pneumococcal-specific IgG antibody response at 28 days PI. Consistent with these observations, TIGR4 inoculated mice displayed significantly increased inflammatory gene expression (IL-1β & IL6) in the lungs and significantly increased levels of circulating inflammatory protein (CCL3) at 7- and 28-days PI respectively.

**Conclusions:** Our mouse model presents a discovery tool to understand the link between acute infections, including pneumonia, and increased cardiovascular disease risk in humans with inflammation as the mechanistic catalyst.

**Graphical Abstract:** 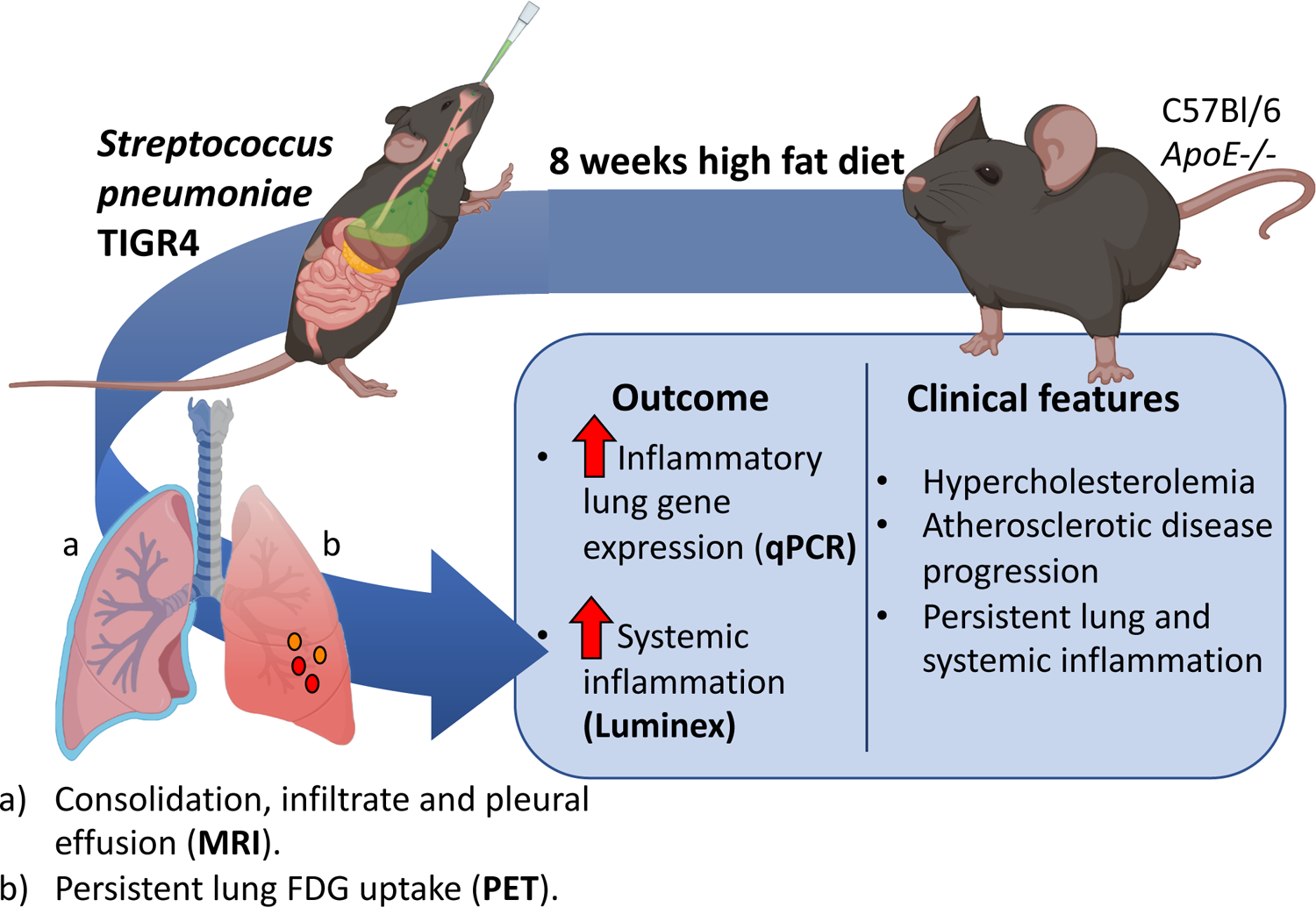

## INTRODUCTION

Patients that survive pneumonia have up to a 4-fold greater risk of cardiovascular disease (CVD) (i.e., myocardial infarction or stroke) in the first 30 days post infection; however, the risk of CVD complications can remain elevated about 2-fold for up to 10 years post-infection(1–4). The mechanisms for this increased CVD risk following community acquired pneumonia (CAP) are still not clear. Dysregulated inflammatory activity related to CAP has been implicated in the progression of atherosclerosis and its CVD complications(1, 5).

Establishing an animal model of pneumonia and atherosclerosis is critical not only for the study of the mechanisms contributing to the increased CVD risk after pneumonia in humans, but also to determine whether atherosclerosis can affect pneumonia risk or progression. For example, it has been suggested that pre-existing atherosclerosis can modify pulmonary immunity or the immunopathogenesis of respiratory infections. Fang et al.(6) showed that hypercholesterolemia induced by 12-16 weeks high fat diet (HFD) in wild-type C57BL/6J mice caused low grade pulmonary inflammation mediated by the activation of the toll-like receptor/nuclear factor kappa-light-chain-enhancer of activated B cells (TLR/NF*κ*B) pathways. Similarly, Ouyang et al.(7) showed that *ApoE-/-* mice on a HFD have increased pulmonary immune cell infiltration at 4 weeks, and increased lung cholesterol content and production of inflammatory mediators at 12 weeks. Compared to wild-type mice, the lungs of *ApoE-/-* mice infected with *Chlamydophilia pneumoniae (C. pneumoniae)* have increased levels of interleukin (IL)-10, IL-6 and IL-4 and reduced cellular infiltration, whereas serum *C. pneumonia*-specific IgG and IgM levels are increased(8).

We conceived our study with the objective of investigating the longitudinal immune responses and changes in lung morphology that occur following induction of pneumonia with *Streptococcus pneumoniae* (*S. pneumoniae)* serotype 4 strain (TIGR4) in *ApoE-/-* mice and monitoring for up to 28 days post-inoculation (PI). We chose *S. pneumoniae* because it is the most prominent cause of CAP in humans and is associated with significant morbidity and mortality worldwide(9). We chose *ApoE-/-* mice because this is an already well-established animal model of pre-clinical atherosclerosis in humans.

## MATERIAL AND METHODS

### Animals

Two groups of *ApoE-/-* male mice aged 6-7 weeks were obtained from the Animal Resources Centre (Figure 1a and b) (West Australia, Australia) and maintained under pathogen-free conditions. Animals were acclimated for 7 days with standard rodent chow before they were transitioned to a HFD containing 21% fat and 0.15% cholesterol to accelerate development of atherosclerosis (Specialty Feeds, Glen Forest, WA, Australia). After 8 weeks of HFD, atherosclerotic plaques were evident in the aortic arch of *ApoE-/-* mice (Supplementary Figure 1a). All animal experiments and procedures were approved by the local ethics committee (Harry Perkins Institute of Medical Research (AE114) and the University of Western Australia (F71731)).

**Figure 1.**
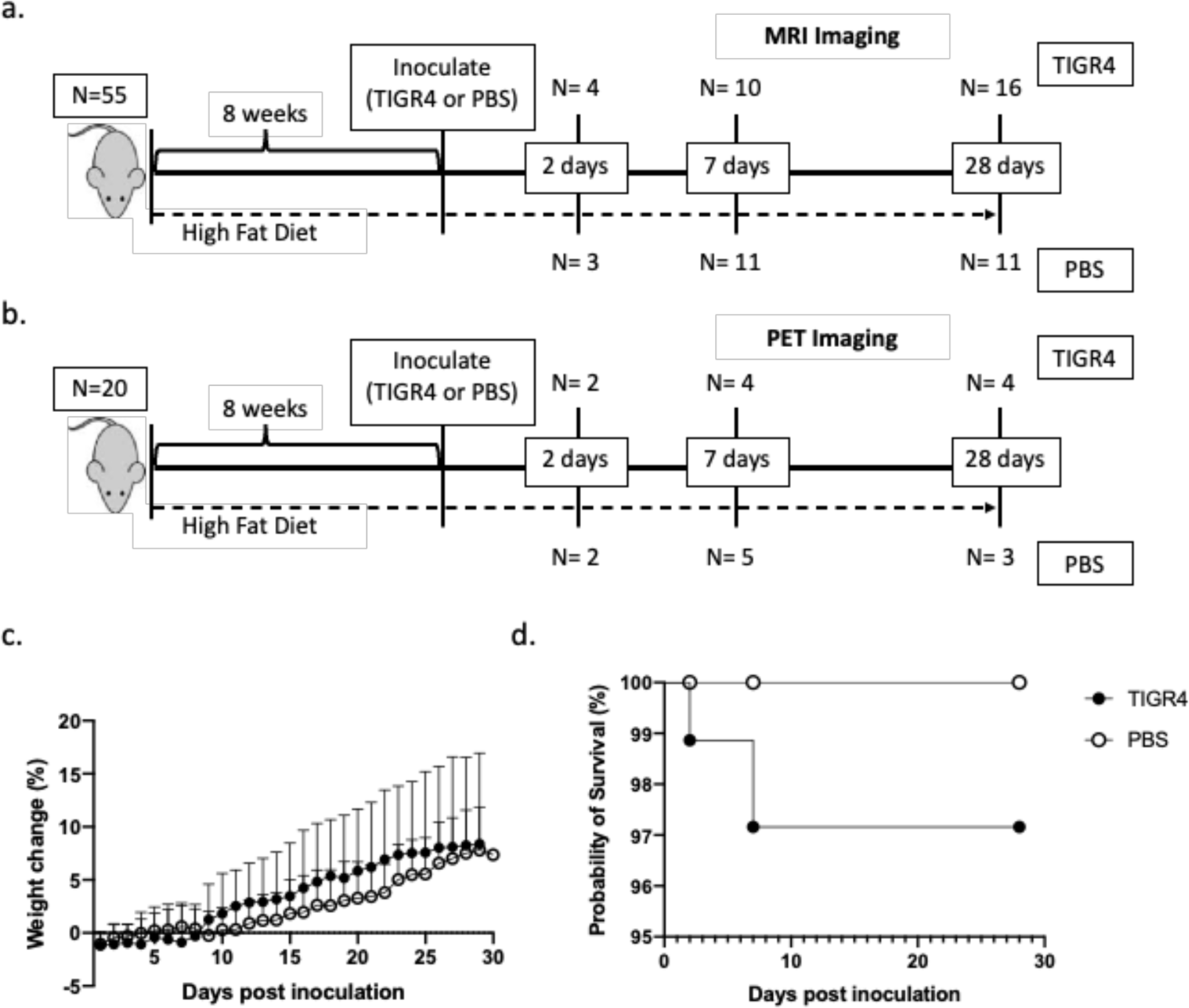
Body weight gain and survival rate. (a) Experimental flow chart. Mice underwent magnetic resonance imaging (MRI) at specific time-points (2, 7 or 28 days PI). Blood was also collected before switch to high fat diet (HFD), 5 weeks into HFD, pre-inoculation and at study end-point ((2, 7 or 28 days post-inoculation (PI)). (b) In parallel, a group of 20 mice were treated the same but assigned for positron emission tomography (PET) imaging. Due to the presence of radiotracer, only blood was collected prior to inoculation and imaging otherwise no tissues were harvested from these mice. (c) Weight loss of mice intranasally inoculated with 10^5^ TIGR4 or PBS. (d) Survival curve of mice intranasally inoculated with TIGR4 or phosphate buffered saline (PBS).

### Bacterial culture

*S. pneumoniae* strain TIGR4 (gifted by Gary Lee, University of Western Australia), was grown overnight on blood agar plates (with 5% sheep blood) or in brain heart infusion broth (SA, ThermoFisher Scientific) until OD_600_=0.4. Aliquots of stock cultures in logarithmic growth were frozen in 10% glycerol (Sigma) and stored at −80°C. Colony forming units (CFU) were determined by plating on blood agar plates and measuring optical density. Bacteria were washed twice and diluted in phosphate buffered saline (PBS) to obtain the appropriate concentrations for the mouse intranasal inoculation.

### Intranasal inoculation

Mice were lightly anesthetized by inhalation with 3% isofluorane (Provet) and inoculated intranasally with 10^5^ CFU of bacteria in a total volume of 40μl. To determine the final dose 3 factors were considered: 1) the dose had to be sufficient to impose weight loss while allowing the mice a chance to recover from the infection, 2) the survival rate had to be greater than 80% and 3) pneumonia had to be detectable by MRI (magnetic resonance imaging) or PET/CT (positron emission tomography/computed tomography) imaging. The final dose was within the range of CFUs from similar mouse studies (10^4^-10^6^ CFU)(10–13), with comparable infection and survival rates and without antibiotic intervention. The dose was initially confirmed in two independent experiments of 10 mice to confirm mortality rate and consistency in MRI imaging. Bacterial inoculation titres were confirmed by serial dilution and plating on blood agar plates. Control mice were challenged intranasally with 40μl PBS. For the final experiment, 75 mice were utilised: 55 mice were assigned for MRI imaging (Figure 1a, TIGR4 n=30, PBS n=25) and the remaining 20 underwent PET imaging (Figure 1b, TIGR4 n=11, PBS=9). Health of mice was monitored twice daily for the first 48 hours post-inoculation and once daily thereafter. Health and disease severity were assessed via weight loss and scoring in any of the following areas: respiration, body posture, eye condition, social interaction, and activity. Based on these criteria and a scoring system developed by the Animal Ethics committee at Harry Perkins Institute of Medical Research, the mice were given a clinical score of healthy [0] to moribund [3]. Any mice displaying a score above 3 were humanely euthanized in accordance with our protocol.

### Bacterial quantification in blood and lungs

Blood was collected from the tail vein, serially diluted in PBS and plated on blood agar plates. To determine bacterial burden in the lungs, supernatants of homogenised lungs were serially diluted in PBS and plated on blood agar plates. Plates were incubated for 18-24 hours at 37°C with 5% CO_2_ and colonies were counted.

### PET/CT imaging

Mice were fasted from food for 4-6 hours (water available). The animals were warmed for 30 minutes (at approximately 30°C) prior to administration of 2-deoxy-2-[fluorine-18]fluoro-D-glucose (^18^F-FDG). Anaesthesia with isoflurane was administered, followed by intravenous (IV) or intraperitoneal (IP) injection of ∼20 MBq of ^18^F-FDG, in a volume no greater than 200μL for IV injection. After completion of one hour uptake, mice were culled, and PET/CT scan was completed on the Bioscan BioPET/CT 105 camera (NSW, Australia). Mice were culled prior to imaging to reduce motion artifact from mouse orientation and movements and to overall improve image quality, and semi-quantitation.

### MRI imaging

Mice were placed under anaesthesia with isoflurane under warming conditions (∼30°C). Once the depth of anaesthesia was adequate, the mouse was moved to the imaging bed where the animal was imaged using the MR Solution MRI 3T scanner. Based on the signal and image quality, a gadolinium-based contrast agent was administered to improve imaging quality if deemed required by the imaging technologist. Anaesthesia with isoflurane was maintained during the scan while respiration and vital signs were monitored remotely. InVivoScope software (Bioscan Inc.) was used for image capture according to the Harry Perkins Cancer Imaging Facility licence.

### Measurement of IgG levels by ELISA

*S. pneumoniae* TIGR4 was cultured in brain heart infusion broth, harvested at mid-log phase and washed twice with PBS. Nunc MaxiSorp 96 well plates (ThermoFisher Scientific) were coated overnight at 4°C with 10^5^ CFU per well of whole TIGR4 pneumococci in 100uL of PBS. Plates were washed three times with 0.05% Tween/PBS and blocked for 1 hour in blocking buffer (PBS with 5% skim milk). Serial 2-fold dilutions of serum in 2% skim milk/PBS were added to the well and incubated for 2 hours at room temperature. After washing, pneumococcal-specific antibodies were detected by incubating with HRP-conjugated goat anti-mouse IgG Ab (ThermoFisher Scientific) diluted 1:10,000 in 2% skim milk/PBS for 1 hour in the dark at room temperature. After washing five times with wash buffer, 3,3’,5,5’-tetramethylbenzidine substrate (Sigma, NSW, Australia) was added and colour development was stopped after 15 minutes by the addition of H_2_SO_4_. Plates were read at 450nm using an Omega plate reader (BMG Labtech, VIC, Australia). The cut-off values were defined using average plus 3 standard deviations.

### Histological analysis

Whole lungs were harvested and embedded in Tissue-Tek OCT (ProScitech, QLD, Australia) and immediately frozen to prevent tissue damage. Samples were stored at −80°C. 10μm sections were cut, air-dried and stained with hematoxylin and eosin (H&E) (Abcam, VIC, Australia). Trichrome staining of the frozen sections was also performed according to the manufacturer’s instructions (Abcam, VIC, Australia) with slight modification. The collagen stain, aniline blue, was replaced with 0.5% fast green in 70% EtOH (Sigma, NSW, Australia). This allowed for red, green and blue colour staining and subsequent colour deconvolution using ImageJ software(14). The sections were mounted in aqueous mounting medium (Fronine). Sections were imaged using a Nikon Eclipse TE2000-U microscope.

### Quantification of gene expression by real-time Polymerase Chain Reaction (PCR)

Ribonucleic acid (RNA) from lungs was extracted using Trizol reagent (ThermoFisher Scientific) and reverse transcribed to complementary deoxyribonucleic acid (cDNA) using the Tetro cDNA Synthesis Kit (Meridian Bioscience, USA) according to the manufacturer’s recommendations. Messenger RNA (mRNA) levels of *gapdh* (Mm99999915_g1), *hprt* (Mm00446968_m1), *il6* (Mm00446190_m1), *tnfa* (Mm00443258_m1), *il1* (Mm00434228_m1), *ifnγ* (Mm00801778_m1), *il10* (Mm00439614_m1), *il17a* (Mm00439618_m1), *il18* (Mm00434225_m1), *nlrp3* (Mm00840904_m1), *ccl2* (Mm00441242_m1), *ccr2* (Mm00438270_01), *smad7* (Mm00484742_m1), *p2×7* (Mm00440582_m1) and *tgfβ* (Mm01178820_m1) were determined by quantitative PCR using Taqman primers and probe (Applied Biosystems, USA) and Taqman Gene Expression Master Mix (Applied Biosystems, USA) on a Rotorgene 6000 (Qiagen). Cycling conditions for TaqMan PCR were 2 minutes at 50°C and 10 minutes at 95°C followed by 45 cycles of 15 seconds at 95°C and 1 minute at 60°C. Samples were performed in triplicate. Data were analysed on the basis of the relative expression method with the formula relative expression 2^-ΔΔCT^, where the amount of the target gene was normalized first to the endogenous reference (HPRT and GAPDH) and then relative to a calibrator (control animal).

### Determination of the serum levels of soluble proteins by multiplex Luminex Assay

Blood was collected from tail vein and centrifuged at 2400rpm for 15 minutes. Serum was collected and stored at −80°C until use. Levels of the following soluble proteins: tumor necrosis factor (TNF)-α, interferon (IFN)-γ, IL-6, IL-1β, IL-5, IL-10, IL-17, CCL3, Dickkopf (Dkk)-1 and matrix metalloproteinase (MMP)-12 were assayed using the multiplex Mouse Magnetic Luminex Assay (R&D Systems, Minneapolis, MN, USA) according to the manufacturer’s instructions. Quantification of proteins were determined using the Luminex 200™ System. The levels of detection for each analyte are: CCL3: 0.45pg/ml, Dkk-1: 31.8pg/ml, IFNγ: 1.85pg/ml, IL-1β: 41.8pg/ml, IL-5: 0.24pg/ml, IL-6: 2.30pg/ml, IL-10: 8.20pg/ml, IL-17: 7.08pg/ml, MMP-12: 0.42pg/ml and TNF-α: 1.47pg/ml.

### STATISTICS

Data are expressed as median (range). All statistical tests were performed with GraphPad Prism 7 (La Jolla, CA, USA). The Kaplan-Meir method was used to compare survival rates. Mann-Whitney was used to compare two groups. Differences with *p*-values <0.05 were considered statistically significant.

## RESULTS

### Intranasal inoculation and mice mortality

To investigate the impact of the selected inoculum of *S. pneumoniae* on the mortality of mice with atherosclerosis, *ApoE^-/-^* mice (n=75) were intranasally inoculated with either 10^5^ CFU of *S. pneumoniae* (n=40) or PBS (control, n=35) after 8 weeks on a HFD (Figure 1a and b). Most TIGR4-inoculated mice developed minor symptoms of infection including ruffled fur, reduced activity, enlarged nasal vestibule, arched backs and squinted eyes during the first 48-72 hours PI. Despite a subtle change in weight loss (Figure 1c), there was no significant evidence of higher mortality (Figure 1d) in infected compared to uninfected *ApoE^-/-^* mice. Three mice died of pneumococcal infection. The first dead mouse died on day 2 PI and cultures of its blood and liver tissue yielded 8.0×10^5^ cfu/ml and 5.3×10^5^ cfu/ml of *S. pneumoniae*, respectively. The second dead mouse died on day 7 PI and while blood culture yielded no bacterial growth, cultures of lung and liver tissues yielded 3.7×10^5^ cfu/ml and 3.3×10^5^ cfu/ml of *S. pneumoniae*, respectively. The third mouse died 2 days PI, yielding 4.3×10^5^ cfu/ml growth in the lung and 3.7×10^5^ cfu/ml growth in the liver. Macroscopic pathological features (e.g. hemorrhagic and mottling) of the lungs and spleen are shown in Supplementary Figures 1b and c. Bacterial growth and morphology are represented in Supplementary Figure 1d. These results confirmed that intranasal inoculation of 10^5^ CFU bacteria produced mild mortality in *ApoE^-/-^* mice, with over 95% survival.

### Intranasal inoculation with S. pneumonia and bacterial burden in tissues and blood

Blood and lung tissues were collected at 2, 7 and 28 days PI. No bacteria were recovered from the lungs and there was no evidence of bacteraemia in infected *ApoE^-/-^* mice that survived until the endpoints of the study. This observation is in line with previous studies of low dose pneumonia in C57Bl/6 mice. Sender *et al.* recovered only 2% of bacteria via bronchoalveolar lavage 6 hours post-intratracheal injection of 10^5^ TIGR4 bacteria in C57Bl/6 mice(15).

### Ongoing inflammation on FDG PET/CT imaging

Evaluation of mouse lung PET/CT scans was performed by a level 3 trained nuclear medicine specialist (SV), who was blinded to the study groups and designations. Representative PET/CT scans are shown in Figure 2. In line with previous PET imaging in pneumonia patients(16), of the 20 mice assigned to PET imaging, all 11 mice that received an intranasal dose of TIGR4 bacteria presented with increased FDG uptake ((standardised uptake value (SUV)) in the lungs compared to PBS inoculated mice (n=8) across all time-points (Figure 3a and b). Of note, at 2- and 7-days PI there was a significant increase (p=0.0025 at 2 days and p=0.0159 at 7 days) in both the average and maximum lung SUVs in TIGR4 inoculated mice compared to PBS. One PBS mouse was excluded from the study due to the presence of a genetic heart defect.

**Figure 2.**
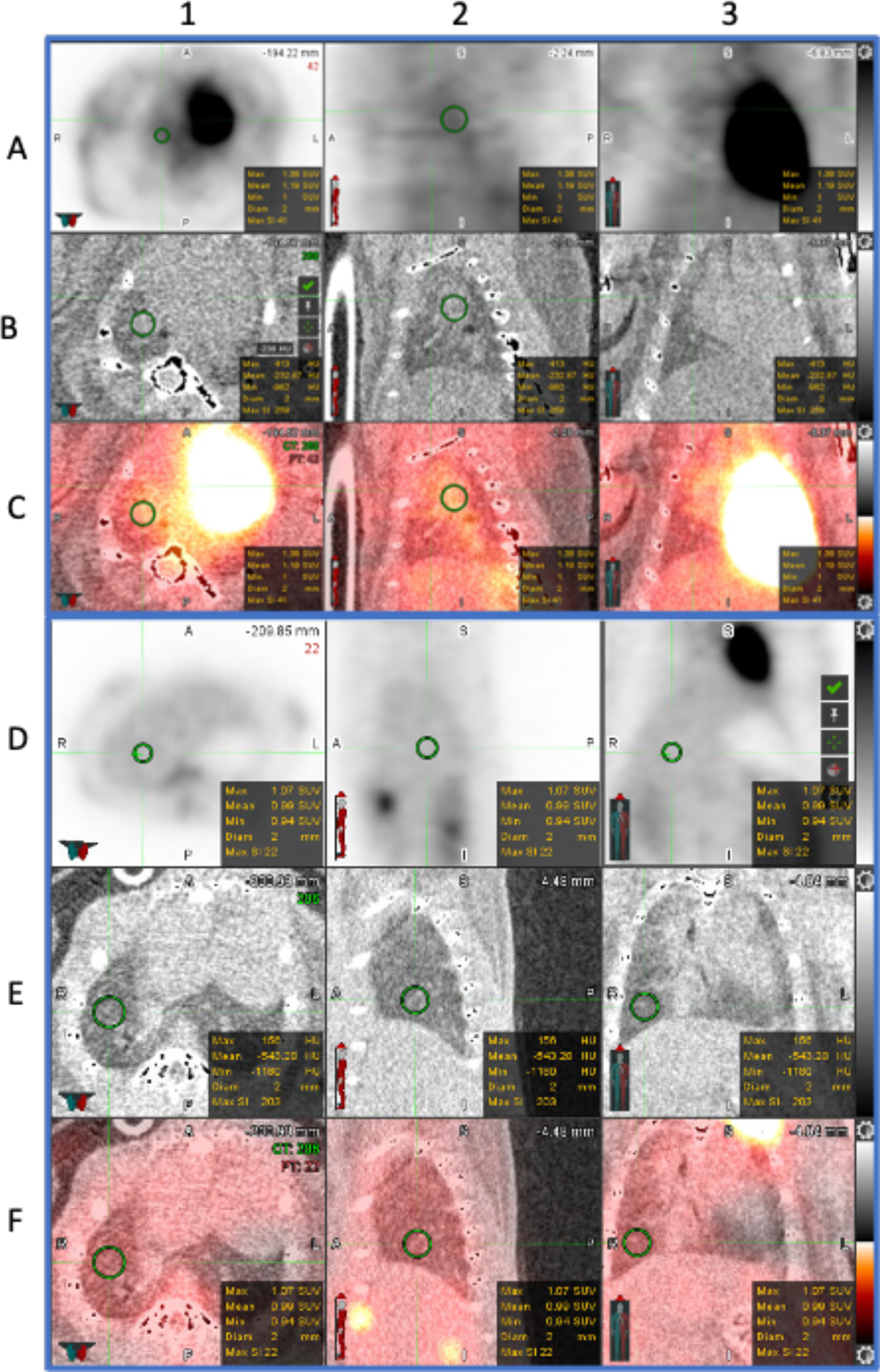
FDG-PET Images from TIGR4 *Streptococcus* infected mice. Rows A-C are representative images from a TIGR4 infected mouse at 28 days post-inoculation (PI). Rows D-F are representative images from a phosphate buffered saline (PBS) inoculated mouse at 28 days PI. Rows A and D are positron emission tomography (PET) scan, B and E are computed tomography (CT) for localisation, C and F are fused PET and CT scans. Regions of interest in row C demonstrate increased fluorodeoxyglucose (FDG) uptake in TIGR4-inoculated mice compared to PBS inoculated mice (row F).

**Figure 3.**
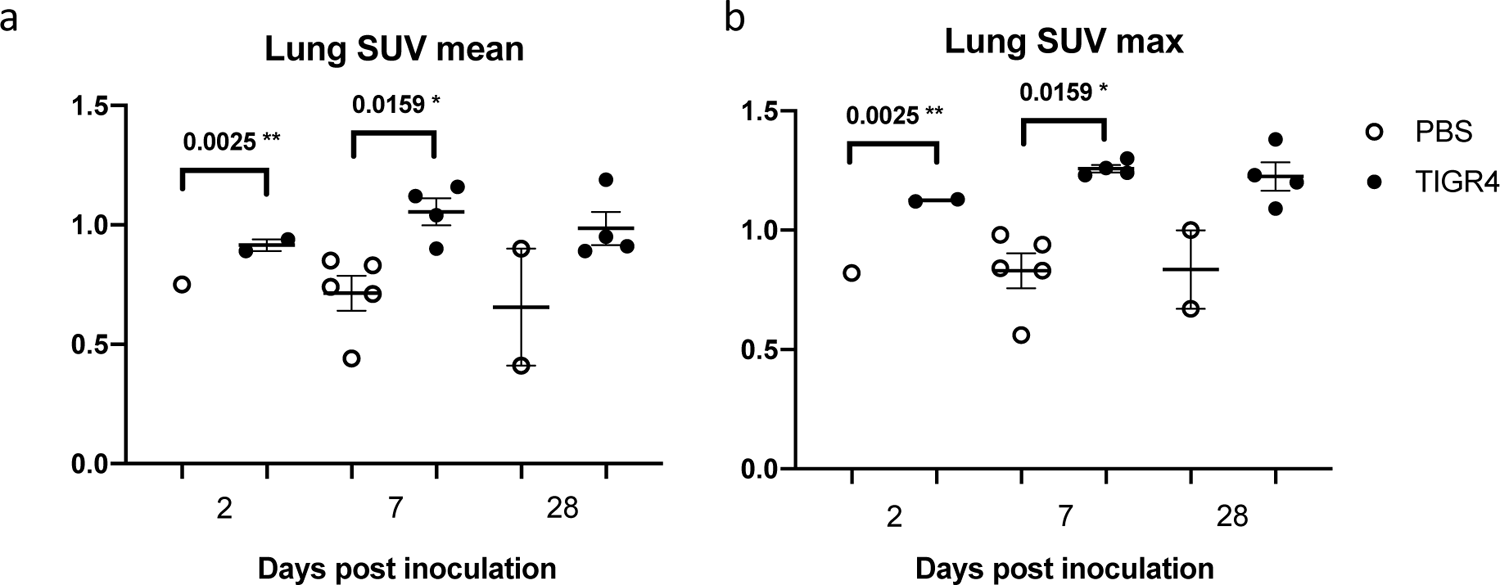
PET imaging of the lungs. Across all time-points there was increased fluorodeoxyglucose (FDG) uptake in the lung of TIGR4 inoculated mice compared to phosphate buffered saline (PBS) inoculated mice. At 2- and 7-days post inoculation there was a significant increase (p=0.0025 ** at 2 days and p=0.0159 * at 7 days) in TIGR4 inoculated mice compared to PBS in both the average and maximum lung standardised uptake values (SUV).

### Intranasal inoculation of *S. pneumoniae* and lung changes on MRI

Evaluation of mouse lungs was performed by a level 3 trained radiologist (TS), who was blinded to the study groups and designations. The 2 mice that succumbed to the TIGR4 infection were not imaged. Of the four TIGR4-inoculated mice that underwent MRI at 2 days PI, three presented radiological findings consistent with lung infection including pleural effusion, consolidation, and various degreess of lung infiltration. 4/8 (50%) TIGR4-inoculated mice scanned at 7 days PI, and 6/15 (40%) TIGR4-inoculated mice scanned at 28 days PI presented with the characteristics described above. None of the PBS-inoculated mice presented with any lung radiological abnormalities that would suggest a respiratory infection. Hence, across all time-points, 13 of the 28 mice inoculated with *S. pneumoniae* presented with radiological findings suggestive of pulmonary infection (Figure 4). The weight the mice that were positive for lung infection were compared to the weights of the remaining TIGR4 mice (Supplementary Figure 3k).

**Figure 4.**
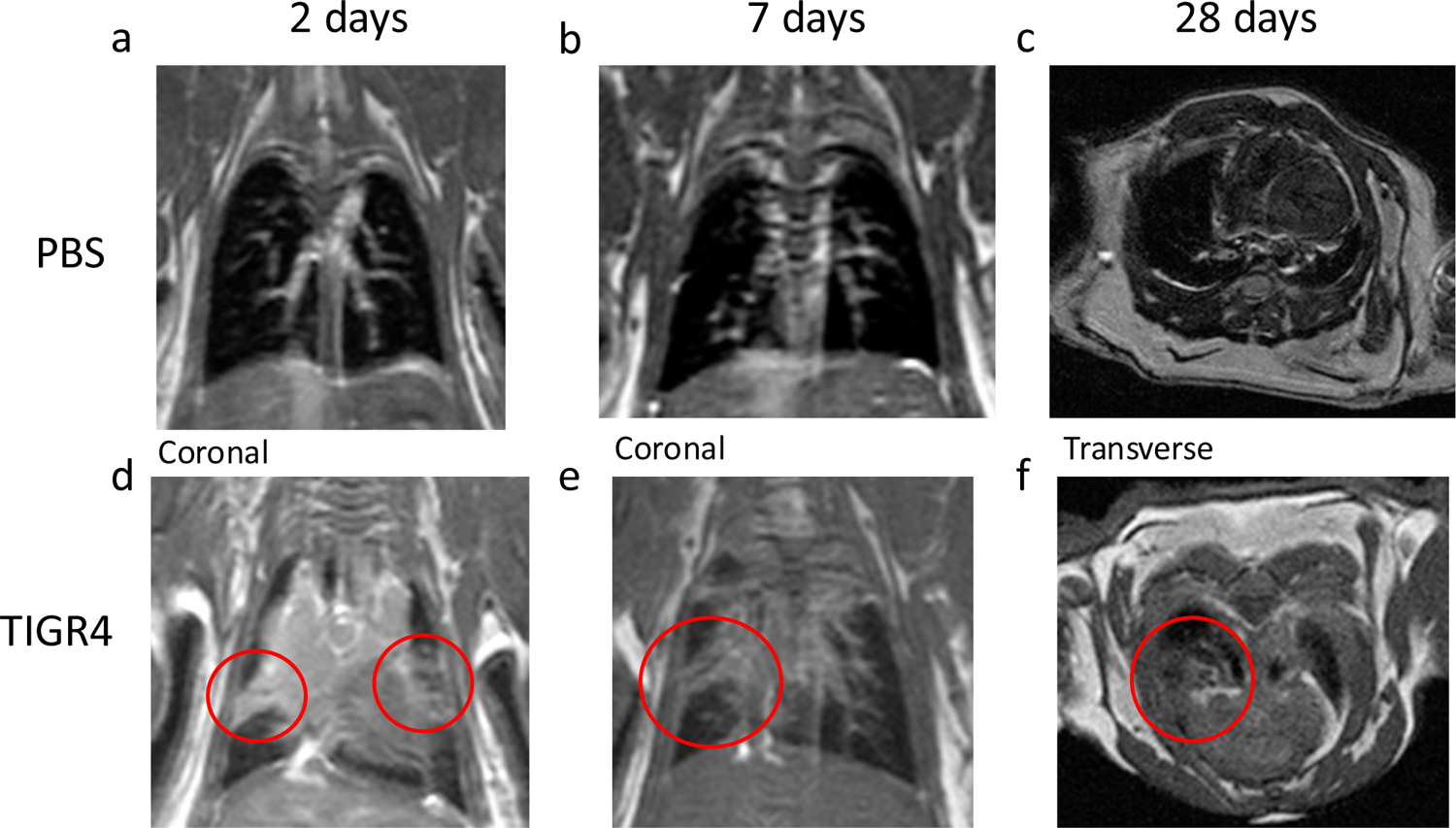
MRI of the lungs. (a) and (d) are representative images of mouse lungs from a phosphate buffered saline (PBS)- and TIGR4-inoculated mouse at 2 days post inoculation (PI). Region of interest (ROI) in (d) demonstrates left and right lung consolidation with infiltration in the TIGR4-inoculated mouse. (b) and (c) are representative images of lungs from a PBS- and TIGR4-inoculated mouse at 7 days PI. ROI highlighted in (e) highlight infiltrate in the right middle lobe. (c) and (f) are representative images of lungs from a PBS- and TIGR4-inoculated mouse at 28 days PI. ROI in (f) demonstrates pleural effusion in the right lung. All scans shown are T1 weighted images.

### Pneumococcal-specific IgG antibody responses

At 2 days PI, one TIGR4-inoculated mouse produced low level of pneumococcal-specific IgG antibody (Figure 5a). No antibody responses were recorded in TIGR4-inoculated mice at 7 days PI (Figure 5b). In contrast, 15/17 (88%) TIGR4-inoculated mice developed an antibody response (Figure 5c, p<0.0001) at 28 days PI. Pneumococcal-specific IgG antibodies were not detected in any of the PBS-inoculated animals. These results are consistent with IgG antibody kinetics to *S. pneumonia* infection(17) and demonstrated that the inoculating dose selected for this study was sufficient to elicit antigen-specific antibody responses.

**Figure 5.**
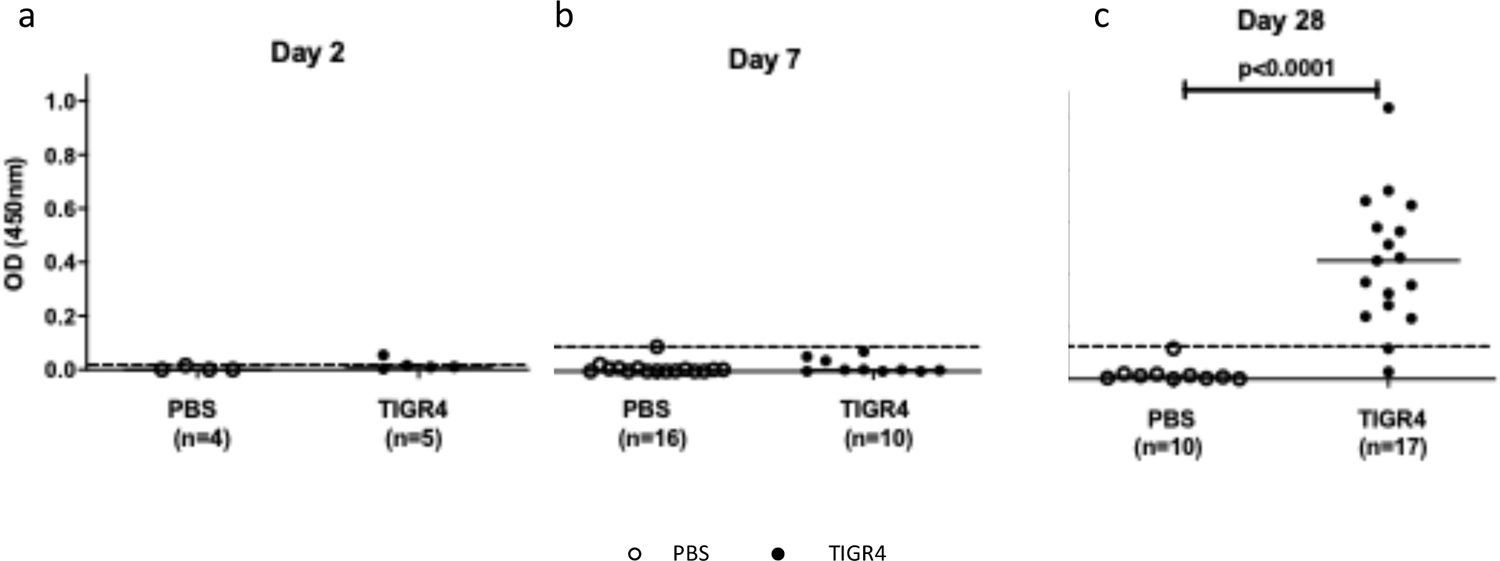
*S.pneumoniae*-specific serum antibody levels. Blood was collected from phosphate buffered saline (PBS) (control) or TIGR4-inoculated mice at specific time-points post-inoculation (PI) (a) is 2 days PI, (b) is 7 days PI and (c) is 28 days PI. Dotted horizontal line represents average + 3x standard deviation. Solid horizontal line represents the cut-off. Solid horizontal line represents the median. OD: optical density

### Lung pathology

Hematoxylin and Eosin (H&E) staining revealed lung remodelling consistent with previous studies in *ApoE-/-* mice(18, 19), however this appears exacerbated in TIGR4-inoculated mice. Progressively, alveolar space and nuclei density increases 2-, 7- and 28-days PI (data not shown), with the greatest difference observed at 28 days PI (Figure 6a). Compared to PBS-inoculated animals, TIGR4-inoculated mice demonstrated increased alveolar septal thickness, suggestive of increased immune cell infiltration. Despite increased lung density and infiltration, there was no significant difference in collagen presence detected in the lungs of TIGR4- and PBS-inoculated mice (Figure 6b).

**Figure 6.**
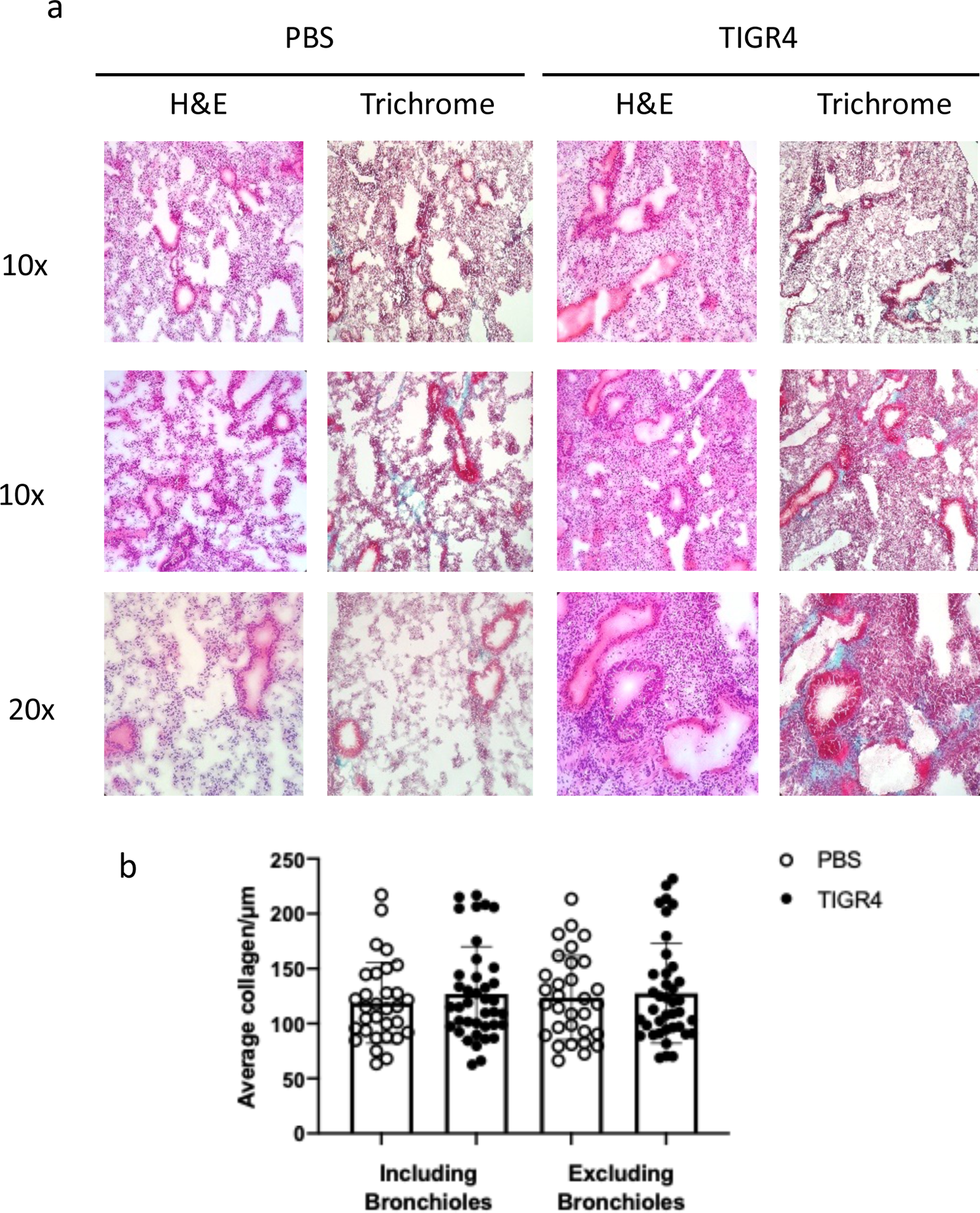
Representative images of lung remodelling in TIGR4- and PBS-inoculated mice at 28 days post-inoculation. (a) Tissue sections were stained with Hematoxylin and Eosin (H&E) or Trichrome. (b) Average lung collagen content per μm in TIGR4- and phosphate buffered saline (PBS)-inoculated mice including and excluding bronchioles.

### Increased IL-1β and IL-6 gene expression in the lungs of *ApoE-/-* mice infected with *S. pneumoniae*

IL-1β gene expression was significantly increased in the lungs of TIGR4-inoculated mice compared to control mice at 7 days PI (p=0.002, Figure 7a). IL-6 gene expression was also significantly higher in TIGR4-inoculated mice at 28 days PI (p=0.01, Figure 7b). There was also a trend for elevated IL-1β (p=0.09) and TNF*α* (p=0.11) gene expression in TIGR4-inoculated mice at 28 days PI (Figures 7a and 7d). In contrast, a trend towards decreased IFN*γ* gene expression (p=0.09) was evident in TIGR4-inoculated mice at 7 days (Figure 7e). There was no difference in TGF-β gene expression in lung tissues between TIGR4- and PBS-inoculated mice at any of the time-points (Figure 7c), confirming the immunohistochemistry collagen results. Gene expression for chemokine receptor (CCR)2, chemokine ligand (CCL)2, IL-10, IL-17, IL-10, NLRP3, purinoceptor 7 (P2X7) and SMAD7 were similar in TIGR4- and PBS-inoculated mice (Supplementary Figures 2a-h). Mice positive for respiratory infection confirmed by MRI were compared to remaining TIGR4 mice, display a trend of increased IL-6, TNF-α, IL-1, NLRP3 and CCL2 at 7 days before decreasing at 28 days PI (Supplementary Figure 3). The results indicate residual local inflammatory dysregulation in mice inoculated with *S.pneumoniae* despite no bacterial burden in the lungs.

**Figure 7.**
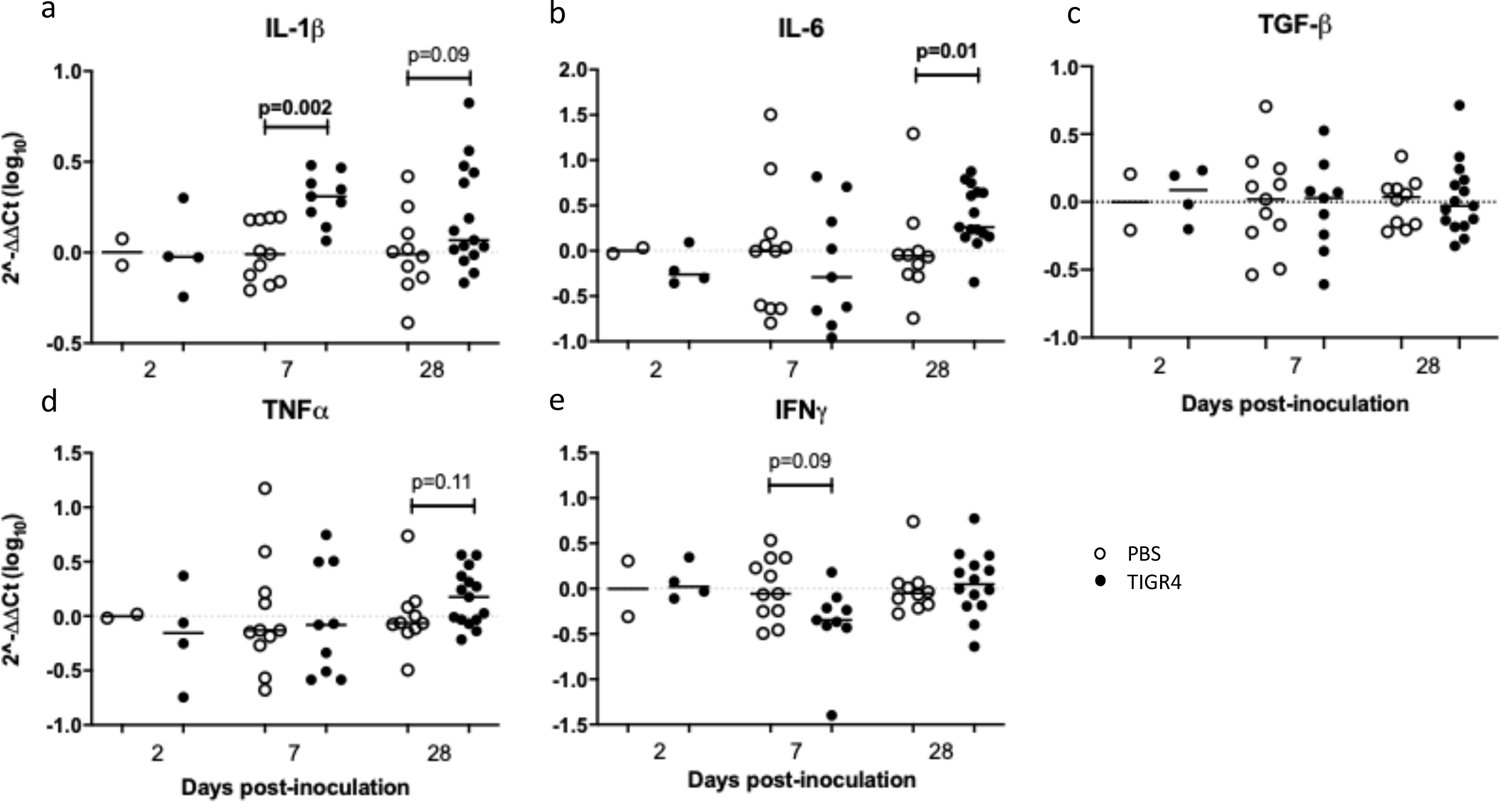
Gene expression of inflammatory mediators in the lungs. Phosphate buffered saline (PBS) control or TIGR4-inoculated mice were sacrificed at specific time-points and ribonucleic acid (RNA) was extracted from lung tissues. mRNA expression levels of (a) interleukin (IL)-1β, (b) IL-6, (c) tumor growth factor (TGF)-β, (d) tumor necrosis factor (TNF)-α and (e) interferon (IF)-γ were quantified by real-time polymerised chain recation (PCR) and normalised against two housekeeping genes (GAPDH and HPRT). Solid horizontal line represents the median.

### Increased circulating levels of IL-6 and CCL3 in *ApoE-/-* mice infected with *S. pneumoniae*

There were no significant differences observed in serum levels for any of soluble proteins at 2 days PI between TIGR4- and PBS-inoculated mice. At 7 days PI, levels of IL-6 were significantly higher in TIGR4-inoculated mice compared to mice inoculated with PBS (Figure 8e, p=0.017). At 28 days PI, CCL3 levels were significantly higher in TIGR4-inoculated mice compared to mice inoculated with PBS (Figure 8b, p=0.007). No differences were observed for levels of Dkk-1 and MMP-12 between TIGR4- and PBS-inoculated mice at 7- and 28-days PI (Figures 8a and h). In most samples, IL-5, IL-1β, IL-10 and IL-17 were below the levels of detection (Figures 8c-d, f-g), whereas TNF-α and IFN-γ were undetectable for all the mice (data not shown). The results indicate ongoing heightened systemic inflammatory activity in mice inoculated with *S.pneumoniae* despite clearance of bacterial infection.

**Figure 8.**
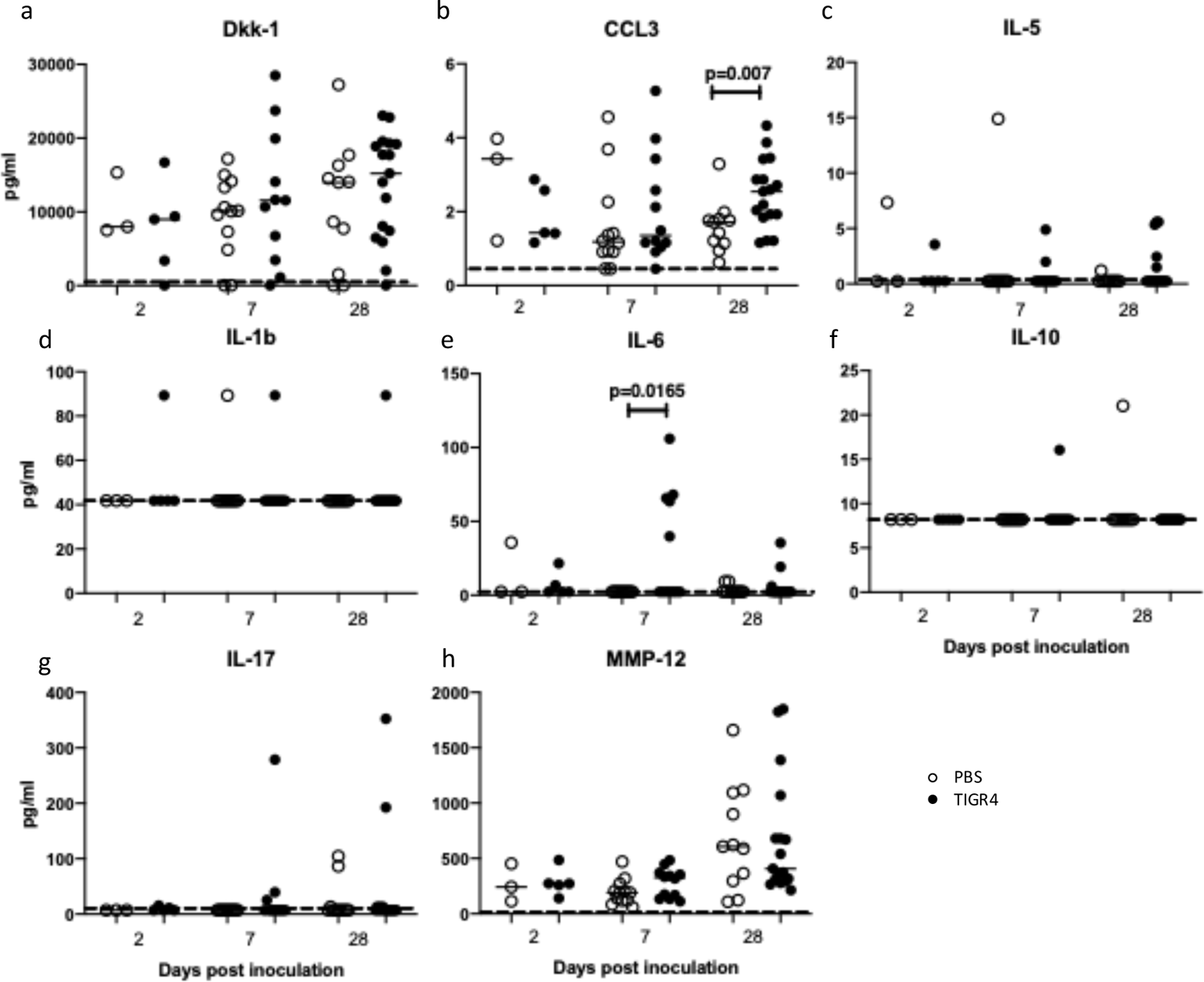
Circulating levels of soluble proteins. Blood was collected from phosphate buffered saline (PBS) control or TIGR4-inoculated mice at specific time-points. Levels of circulating (a) dickkopf-related protein 1, (b) chemokine ligand (CCL) 3 (also known as macrophage inflammatory protein 1-α, (c) interleukin (IL)-5, (d) IL-1β, (e) IL-6, (f) IL-10, (g) IL-17, and (h) matrix metalloproteinase (MMP) 12 were assessed. Horizontal line represents the median. Dotted line represents minimum level of detection of the Luminex assay. N values: 2d: Inf= 5 Control= 3. 7d: Inf= 12 Control= 12. 28d: Inf= 17 Control=11.

## DISCUSSION

To date, few alternative multiple comorbidities animal models of acute infection and atherosclerosis have been described(10,20–22). In contrast to our model, these other models utilise viral pathogens (e.g., influenza virus(21)), alternative routes of infection (eg: intraperitoneal) that are more relevant to sepsis than pneumonia(22), intervene with antibiotics to improve mortality(10), or do not describe the lung pathology associated with the infectious challenge(10, 20). Our model utilises a low infectious dose to establish infection with *S. pneumoniae*, the most important bacterial cause of pneumonia in humans(23–25), via intranasal inoculation; hence, being representative of the pneumococcal infection that is acquired via the respiratory route in humans. Additionally, our model produces a range of pathology that includes no detectable illness, subclinical disease, clinical infection, and infection-induced death thus resembling that full scope of pathology produced by exposure to respiratory pathogens in humans. Finally, by foregoing antibiotics, we do not introduce any putative additional confounder from such bioactive compounds (for example, some antibiotics such as macrolides, tetracyclines and fluoroquinolones can exert important anti-inflammatory activity)(26–28). Hence, to the best of our knowledge, this is the first study to establish a longitudinal model of *S.pneumoniae* infection with a wide range of pulmonary and systemic pathology in *ApoE-/-* mice beyond the acute phase of the infection and well into the clinical recovery of surviving animals.

It has been proposed that the introduction of a respiratory infection leads to a mounting synergistic inflammatory response that could lead to adverse cardiovascular events. In CAP, exposure of alveolar epithelial cells and resident macrophages to *S. pneumoniae* stimulates the production of inflammatory cytokines such as IL-1β, IL-6 and TNF*α*, as well as chemokines(25,29,30). The duration and magnitude of the inflammatory response have been linked to disease severity and clinical outcome. In a longitudinal study of 247 hospitalised patients with CAP, cytokine and chemokine levels were highest at hospital admission and while most decreased at 6 weeks follow-up, some markers remained elevated(31), suggesting ongoing residual inflammation. This was confirmed in a recent study of 22 CAP survivors, with 68% of subjects displaying increased ^18^FDG uptake in the lungs, 30 to 45 days after their hospital discharge(16).

In agreement with these observations, TIGR4-inoculated mice display delayed elevated inflammatory response in the lungs (e.g. IL-6 at 28 days PI) and systemically (e.g. CCL3 at 28 days PI). Interestingly, systemic levels of IL-6 peaks at 7 days PI. IL-6 plays an important role in linking innate to an acquired immune response by promoting differentiation of naïve CD4+ T cells(32). IL-6 can serve as an activator for other pro-inflammatory cytokines including IL-1 and TNF-α(33).

In response to an infection, IL-6 stimulates a range of signalling pathways including NF-κB, enhancing the transcription of the mRNA of inflammatory cytokines including IL-6, TNF-α, and IL-1β. TNF-α and IL-1β in turn also activate transcription factors to produce more IL-6(34).

A study published by Bacci *et al.* correlate increased IL-6 and TNF-α with worse outcomes in pneumonia patients(33). While TNF-α and IL-6 gene expression in the lung from TIGR4-inoculated mice at 28 days was not statistically significant, results do appear to show an upward trend. Patients discharged with pneumonia leave hospital with ongoing subclinical inflammation(35), and in an *ApoE-/-* model of bacterial tuberculosis(36), there is a delayed adaptive immune response. Therefore, it’s possible in our study that not enough time PI had lapsed and a later time-point may be worth investigating to elucidate long-term inflammatory changes. Neutrophils play a key role in response to controlling a pneumococcal pneumonia infection(37).

CCL3 acts as a chemotactic factor for different leukocyte subsets and has also been shown to increase during development of atherosclerotic lesions(38, 39). It has been shown that during acute inflammation triggered by lipopolysaccharide stimulation, CCL3 induces chemotaxis for neutrophils to atherosclerotic lesions(39). Our data supports the induction of CCL3 in response to an acute infection and atherosclerosis disease progression (Figure 8b). Altogether there is a lot of overlap in immune response to atherosclerosis and pneumonia, sharing a lot of the same immune pathways that could be pathologically relevant for atherosclerotic progression.

Previously associated with lung fibrosis, epithelial cell proliferation, acute lung inflammation, increased atherosclerotic apoptosis and enlarged and destabilized plaques, Dkk-1 has been identified in both a respiratory infection and atherosclerosis setting(40, 41), (42). Due to the majority of Dkk-1 being platelet produced and its involvement in atherosclerosis, it has been recently suggested as an independent risk factor for major adverse cardiovascular events(42). Progressively increasing amounts of Dkk-1 were measured in TIGR4 inoculated mice compared to PBS (Figure 8a).

In terms of lung fibrosis, HFD and deletion of the *ApoE* gene can induce morphological changes to the lungs including lipidosis, increased pulmonary inflammation and increase in alveolar septal thickness(7,19,43). These are consistent with findings in this model, with altered lung morphology in PBS-inoculated mice in response to hypercholesterolemia, and TIGR4-inoculated mice presenting with more severe remodelling as a result of inflammatory infiltrate caused by TIGR4-inoculation. The lungs of TIGR4 inoculated mice have increased nuclei and inflammatory gene expression indicating an elevated sublethal amount of inflammation. While no difference in systemic Dkk-1 or collagen was detected, it could be argued that not enough time PI had elapsed to see the changes observed in patients. Patients have been documented to be at 2-8-fold increased risk of adverse events for up to 10 years after hospital discharge for pneumonia infection(1). Moreover, the novel Dkk-1 biomarker requires further investigation to validate its clinical relevance in pneumonia and role in prediction of adverse events.

There are some limitations associated with our study. Firstly, we did not investigate the cellular immune responses in the lungs that may shed light on the lack of recovered bacteria/bacterial resistance. However, as we administered a low dose of TIGR4 bacteria in consideration of a low mortality rate, it is possible that the mice experienced a more acute case of pneumonia and therefore a low-level of pneumonia infection. In this scenario, it was also predicted that all observations made would be applicable and to increase in line with a higher dose. Moreover, the significant increase in pneumococcal-specific antibody response observed at 28 days PI is in line with the strong systemic CCL3 response indicative of exposure to *S.pneumoniae*. Alternatively, it is possible that the pre-existing inflammatory response activated by atherosclerosis was enough to clear the low intranasal dose of *S.pneumoniae.* However, this premise will require further investigation. The TIGR4 strain used in our studies is an invasive strain that has been shown to cause pneumonia and lethal systemic disease following intranasal challenge(10–12,44). This strain has been associated with higher rates of pneumonia and enhanced disease severity in C57BL/6 mice, the genetic background of the *ApoE-/-* mice used in the current study compared to BALB/c mice(44).

## CONCLUSIONS

In summary, our animal model utilising *S.pneumoniae* and *ApoE-/-* mice on a HFD offers a more accurate representation of pneumonia disease in humans with pre-existing atherosclerotic disease. Hence, our model is relevant to the study of the changes that could occur following an episode of pneumonia on subclinical atherosclerosis and future cardiovascular events. The suggested underlying mechanism for plaque rupture and adverse events in patients with pneumonia is persistent inflammation from the respiratory infection(1,3,45), (46). Our study suggests that a delayed and robust pro-inflammatory response to *S. pneumoniae* has the potential to contribute to the increased risk of cardiovascular complications that is seen among pneumonia survivors(35, 47).

## ABBREVIATION LIST

CAP: community acquired pneumonia

CCR: chemokine receptor

CCL: chemokine ligand

CFU: colony forming units

CVD: cardiovascular disease

18F-FDG: 2-deoxy-2-[fluorine-18]fluoro-D-glucose

H&E: Hematoxylin and Eosin

HFD: high fat diet

IF: interferon

IL: interleukin

IP: intraperitoneal

IV: intravenous

MMP: matrix metalloproteinase

MRI: magnetic resonance imaging

PBS: phosphate buffered saline

PCR: polymerase chain reaction

PET/CT: positron emission tomography/computed tomography

PI: post inoculation

RNA: ribonucleic acid

SUV: standardised uptake value

TNF: tumor necrosis factor

## ACKNOWLEDGEMENTS

The authors gratefully acknowledge Lincoln Codd, Brenton O’Mara, Kirsty Richardson, Liesl Celliers, Penny Maton and Roslyn Francis for technical assistance. The authors acknowledge the facilities, and the scientific and technical assistance of Microscopy Australia at the Centre for Microscopy, Characterisation & Analysis, The Cancer Imaging Facility at Harry Perkins Institute of Medical Research, The University of Western Australia, a facility funded by the University, State and Commonwealth Governments.

## ETHICS

All animal procedures were carried out in accordance with both the Western Australian Animal Welfare Act and National Institute of Health guidelines. This project and its procedures were approved by the animal ethics committee of the Harry Perkins Institute for Medical Research (AE114) and the University of Western Australia (F71731).

---

**Supplementary Figure 1. Characteristics of TIGR4-inoculated *ApoE-/-* mice.** (a) Representative images of atherosclerotic plaques in the aortic arch of *ApoE-/-* mice after 8 weeks high fat diet. (b) Morphological changes in the lungs of mice inoculated with TIGR4 bacteria. (c) Morphological abnormalities of black and white patches on the spleen of TIGR4 inoculated mice. (d) *Streptococcus pneumoniae* serotype 4 (TIGR4) growth on blood agar plate.

**Supplementary Figure 2. Gene expression of inflammatory mediators in the lungs.** Phosphate buffered saline (PBS) control or TIGR4-inoculated mice were sacrificed at specific time-points and ribonucleic acid (RNA) was extracted from lung tissues. mRNA expression levels of (a) C-C chemokine receptor type (CCR) 2, (b) C-C motif chemokine ligand (CCL) 2, (c) NLR family pyrin domain containing 3 (NLPR3), (d) P2X purinoceptor 7 (P2X7), (e) SMAD family member 7, (f) interleukin (IL)-17, (g) IL-18 and IL-10 were quantified by real-time polymerase chain reaction and normalised against two housekeeping genes (GAPDH and HPRT). Solid horizontal line represents the median.

**Supplementary Figure 3. Gene expression of inflammatory mediators in lungs from TIGR4-inoculated animals stratified based on radiological findings.** TIGR4-inoculated mice with evidence of respiratory infection confirmed by magnetic resonance imaging (MRI) were compared to mice with no radiological abnormalities at respective timepoints (2, 7 and 28 days PI). Ribonucleic acid (mRNA) expression levels of (a) interleukin (IL)-6, (b) tumor necrosis factor (TNF)-α, (c) IL-1, (d) interferon (IF)-γ, (e) IL-10, (f) IL-17, (g) NLR family pyrin domain containing 3 (NLPR3), (h) IL-10, (i) C-C motif chemokine ligand 2 (CCL) and (j) P2X purinoceptor 7 (P2X7) were quantified by real-time polymerase chain reaction and normalised against two housekeeping genes (GAPDH and HPRT). Solid horizontal line represents the median. (k) Weight loss of TIGR4-inoculated mice stratified according to the absence or presence of respiratory infection confirmed by MRI.

