## Supplementary figures 1-3 for "A multiple comorbidities mouse model to assess atherosclerosis progression following lung infection in *ApoE* deficient mice"

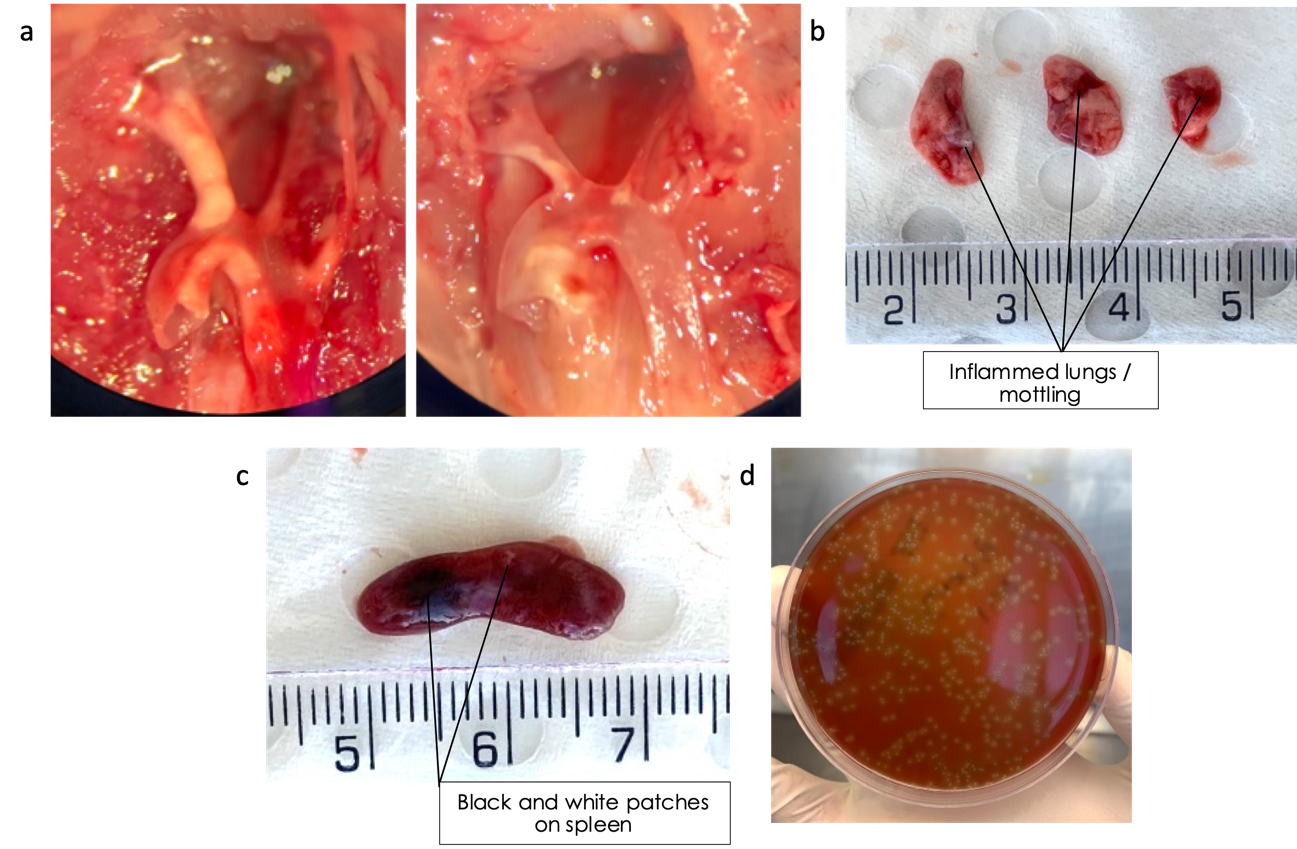

**Supplementary Figure 1. Characteristics of TIGR4-inoculated *ApoE-/-* mice.** (a) Representative images of atherosclerotic plaques in the aortic arch of *ApoE-/-* mice after 8 weeks high fat diet. (b) Morphological changes in the lungs of mice inoculated with TIGR4 bacteria. (c) Morphological abnormalities of black and white patches on the spleen of TIGR4 inoculated mice. (d) *Streptococcus pneumoniae* serotype 4 (TIGR4) growth on blood agar plate.

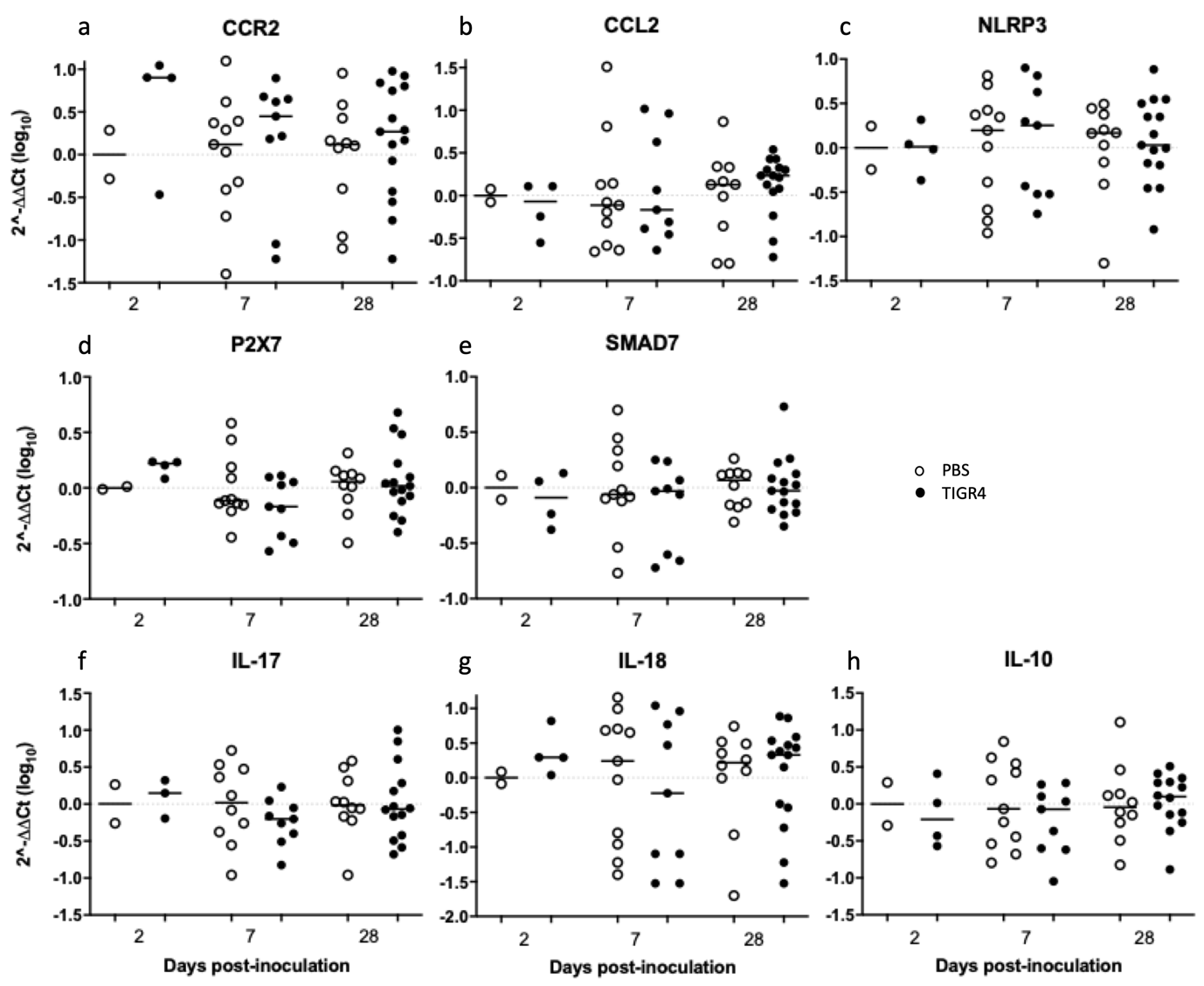

**Supplementary Figure 2. Gene expression of inflammatory mediators in the lungs.** PBS control or TIGR4-inoculated mice were sacrificed at specific time-points and RNA was extracted from lung tissues. mRNA expression levels of (a) C-C chemokine receptor type 2, (b) C-C motif chemokine ligand 2, (c) NLR family pyrin doman containing 3, (d) P2X purinoceptor 7, (e) SMAD family member 7, (f) interleukin (IL)-17, (g) IL-18 and IL-10 were quantified by real-time PCR and normalised against two housekeeping genes (GAPDH and HPRT). Solid horizontal line represents the median.

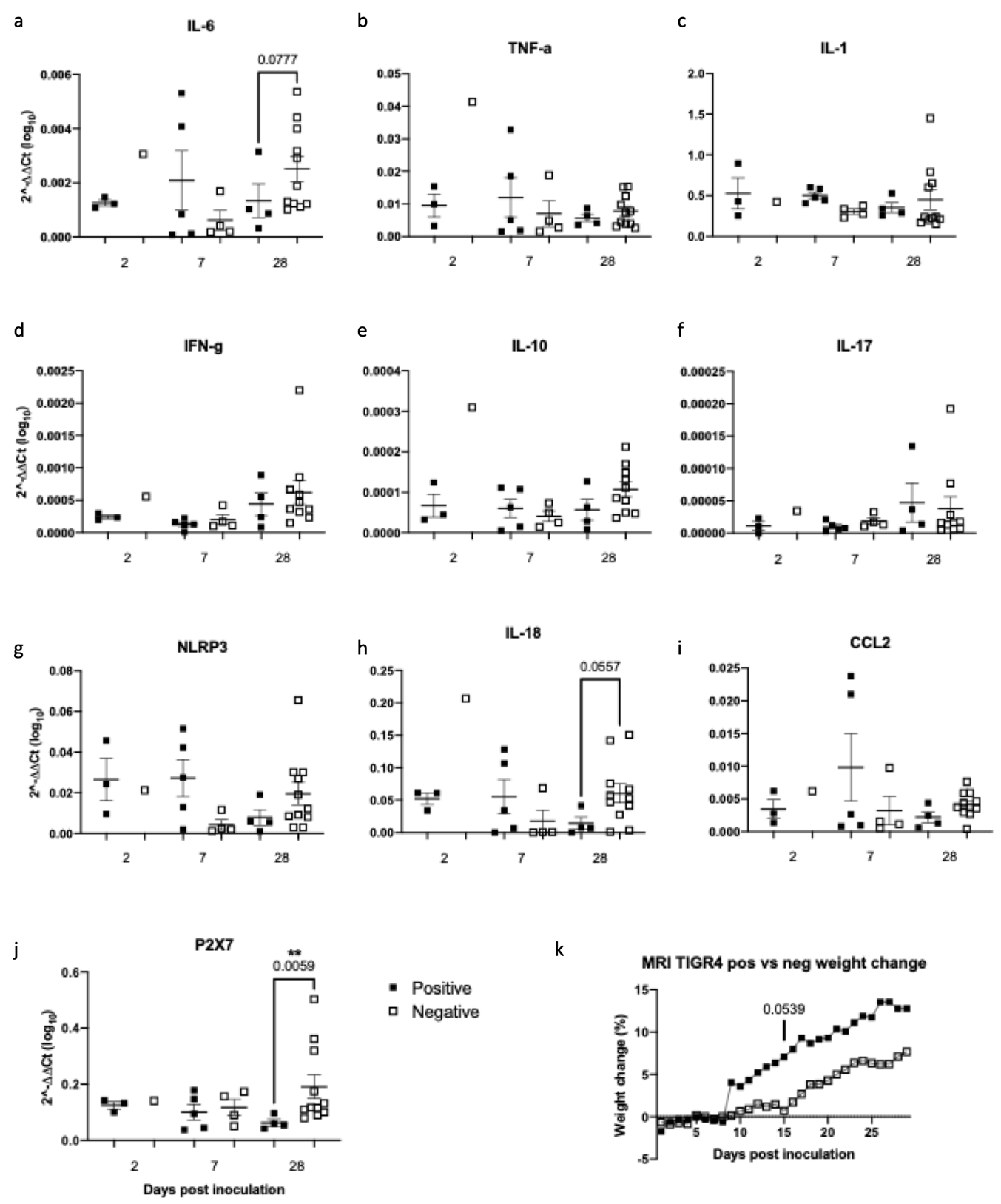

**Supplementary Figure 3. Gene expression of inflammatory mediators in lungs from TIGR4-inoculated animals stratified based on radiological findings.** TIGR4-inoculated mice with evidence of respiratory infection confirmed by MRI were compared to mice with no radiological abnormalities at respective timepoints (2, 7 and 28 days PI). mRNA expression levels of (a) interleukin (IL)-6, (b) tumor necrosis factor-α, (c) IL-1, (d) interferon-γ, (e) IL-10, (f) IL-17, (g) NLR family pyrin domain containing 3, (h) IL-10, (i) C-C motif chemokine ligand 2 and (j) P2X purinoceptor 7 were quantified by real-time PCR and normalised against two housekeeping genes (GAPDH and HPRT). Solid horizontal line represents the median. (k) Weight loss of TIGR4-inoculated mice stratified according to the absence or presence of respiratory infection confirmed by MRI.
